# Harnessing light heterogeneity to optimise controlled environment agriculture

**DOI:** 10.1101/2024.08.20.608762

**Authors:** Will Claydon, Ethan J. Redmond, Gina YW Vong, Alana Kluczkovski, Alice Thomas, Phoebe Sutton, Katherine Denby, Daphne Ezer

## Abstract

Yield is impacted by the environmental conditions that plants are exposed to. Controlled environmental agriculture provides growers with an opportunity to fine-tune environmental conditions for optimising yield and crop quality. However, space and time constraints will limit the number of experimental conditions that can be tested, which will in turn limit the resolution to which environmental conditions can be optimised. Here we present an innovative experimental approach that utilises the existing heterogeneity in light quantity and quality across a vertical farm to evaluate hundreds of environmental conditions concurrently. It proposes a three-phase workflow for identifying critical light variables, which can guide targeted improvements in yield and energy use. Using an observational study design, we identify features in light quality that are most predictive of biomass in different microgreens crops (kale, radish and sunflower) that may inform future iterations of lighting technology development for vertical farms. The findings suggest that light quality, rather than just light intensity, plays a crucial role in uniform crop yields and that light sensitivities are variety-specific, highlighting the importance of tailored light recipes for different crops.

## Introduction

Plants are sensitive to minor changes in environmental conditions, such as light, temperature and humidity. For instance, Arabidopsis is sensitive to as little as a 2°C change in temperature (1). Controlled environmental agriculture practices, such as vertical farming, allow growers to customise the growing conditions of crops to maximise yield for each species (2). Minor environmental differences may also impact agriculturally-important qualities other than yield, such as aesthetic qualities, nutrient concentration and taste compounds (3–5). Growers need to balance these requirements when designing their optimised vertical farm conditions.

To optimise vertical farm conditions, researchers will usually employ a complete randomised experimental design in which they measure crop traits in several different conditions, a strategy that has been successfully deployed in many studies to optimise light, temperature and humidity separately (6–8). However, these studies do not consider how changing one growth condition may impact another. Other researchers have utilised multifactorial experimental designs, especially full factorial designs, to combinatorially assess the impact of several environmental variables concurrently (9–11). For instance, a 4×4×3 full factorial design was used to find optimal combinations of root temperature, air temperature, and light intensity for lettuce growth (8). In contrast, other groups wished to optimise across 6 environmental parameters at 3 levels, which would have required 729 experimental treatments using a full factorial design, but they were able to select 27 combinations of conditions to test by using the Taguchi Method to generate an Orthogonal Array of treatment combinations (2). However, even the Taguchi method requires a relatively large number of treatment conditions, which would be difficult to implement in a small vertical farm. Moreover, any randomised experimental design will assume that treatments are homogenous within each treatment block, when in fact light, temperature and humidity may vary within the physical space due to the layout of lights and the air flow, among other factors, microclimatic conditions that have been modelled in digital twins of vertical farms (12,13).

Although the physiological impact of heterogeneous microclimates in vertical farms has not been fully investigated, it has been widely established that in traditional field-based agriculture, conditions are not uniform within a field, resulting in heterogeneity in crops. Soil topography can vary across and within fields, which has shown to impact how soil can retain water and by proxy impact yield (14). It has also been shown that increasing the distance from the edge of a field that a crop is grown in can increase yield (15). Moreover, crop yields can also be impacted by the landscape that surrounds a field and the amount of shading that a crop receives (16). Environmental variation within the field is detrimental to both food security and to profits made by farmers. This is because within field variation impacts how a crop is shaped, its size, colour and yield, all factors that can lead to crops failing quality checks and being disposed of (17). Vertical farming removes sources of heterogeneity such as soil topography and landscape. However, there are still sources of heterogeneity that occur within vertical farms as shown here.

In this paper, we have taken advantage of the innate heterogeneity of light intensity and quality that exists in a vertical farm to suggest an innovative observational experimental design for condition optimisation. We measured the position-specific light intensity and quality across 256 different positions in a vertical farm and modelled how these factors impact biomass of different microgreen crops. We suggest that exploiting existing heterogeneity in vertical farms to perform observational studies will enable us to optimise treatments using less space-intensive experimental set-ups than in traditional environmental optimization approaches, allowing researchers to test hundreds of different combinations of conditions in a single experiment.

## Materials and methods

### Measuring light heterogeneity in a vertical farm

This work was performed in the Grow It York vertical farm (located in York city centre, UK) which uses LettUs Grow (Bristol, UK) aeroponic technology. Plants were grown in 2500 cm^2^ trays, with four trays per bed, with four beds stacked vertically, totalling 16 trays. Above each bed was a set of three LED lighting fixtures, Horti-blade BRWFR-4 spectrum, from Vertically Urban. Due to space constraints during retrofitting, the LED fixtures were positioned unequally above the bed with two fixtures towards the front and one towards the back.

To establish the extent of light heterogeneity within the vertical farm, we divided each tray into 16 unique sections each with an area of 83.7 cm^2^. Microgreens were also grown around the edge of the tray (9 cm width at the left and right sides of the mat and 11.5 cm at the top and bottom) surrounding the 16 squares in the centre of the mat. The plants in these edge areas were not measured as part of this experiment to negate edge effects in our samples (see **Figure S1**). Each position was given a unique x and y coordinate and in these same positions, plants were grown and biomass recorded. Across all beds this resulted in 256 unique sampling positions. Both Photosynthetically Active Radiation (PAR) and spectral irradiance were measured once in these positions at the level of the tray (at the plant growing level) in the absence of plants within a month of the experiment taking place. PAR was measured using a PAR Special from Skye industries. Spectral irradiance was measured using an Ocean Fire Spectrometer from Ocean View. which could then be used to calculate the mean blue (B), red (R), and far-red (FR) light intensity in each position

Plants were grown under a spectrum consisting of 14% **Blue** 14% **Green** 64% **Red** 7% **Far-Red** light, characterised by blue (400-500 nm), green (500-600 nm), red (600-700 nm), and far-red wavelengths (700-780 nm). For our experimental purposes, we measured narrower waveband ranges: 445-456 nm for blue, 653-668 nm for red and 725-735 nm for far-red (18,19), which correspond to plant photobiological sensitivity, and the LED emission peaks of the Vertically Urban lighting system (**Figure S2**). For the remainder of the manuscript, when we refer to ‘ blue’, ‘ red’ and ‘ far-red’, we refer to the average light intensities in these narrower wavebands. The mean intensities were calculated for each experimental waveband at each of the 256 positions.

### Observational Experiment

All plants were grown under 12 hours of light per day which delivered a PAR reading in the range of 263-568 µmol m^−2^ s^−1^, resulting in 37.99-79.36 µW/cm^2^/nm of blue light, 54.75-62.32 µW/cm^2^/nm of red light and 5.43-14.43 µW/cm^2^/nm of far-red light (Table S1). These light values are expressed as ranges due to the light heterogeneity in the facility (**Table S2)**. Microgreens were grown on Growfelt (Whitworth, UK) wool carpet matting. An aeroponic system was used maintaining the pH at 5.9 and electrical conductivity at 1.7 with Hydromax grow A and B nutrient solutions. The aeroponic system was activated for one minute every four minutes. Each microgreen variety, as outlined in **Table 1**, was grown in one tray per bed in the arrangement shown in **Figure S3**. The beds are vertically stacked in the facility. This equalled four trays per variety giving 64 unique measurement positions per microgreen variety. All samples were grown for the period specified in **Table 1**. Each unique position was harvested individually and fresh biomass recorded.

**Table 1:** Description of microgreen varieties and harvest times.

| Species | Common name | Name in text | Germination Time (days) | Time to harvest after germination (days) |
| --- | --- | --- | --- | --- |
| *Brassica oleracea* | Kale (Dwarf blue) | Kale 1 | 4 | 7 |
| *Brassica oleracea* | Kale (Dwarf green curled) | Kale 2 | 4 | 7 |
| *Helianthus annuus* | Sunflower (Black Seeded) | Sunflower | 3 | 7 |
| *Raphanus sativus* | Radish (Sangria) | Radish | 4 | 5 |

### Generating mixed effects models

Our aim was to predict biomass as this is the yield metric for microgreens. We used a mixed effects model to predict biomass per position (g/83.7 cm^2^), using the lme4 package in R (v4.3.2) (20,21). Our assumptions were that bed and position within a tray would both be confounding factors influencing biomass. We also assumed that the impact of bed and position within a tray would depend on the species/variety of microgreen. Our baseline model was as follows, where m is the microgreen species/variety (**Table 1**), b is the bed, and c is the position within the tray (i.e. centre, edge or corner, **Figure S1**).

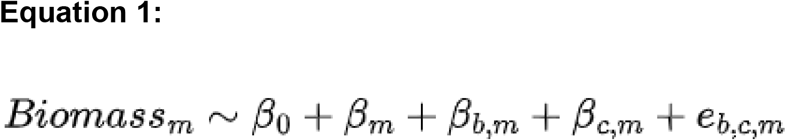

Next, we added light intensity (PAR, µmol m^-2^ s^-1^) to the baseline model, allowing for light intensity to have a different effect on each microgreen variety:

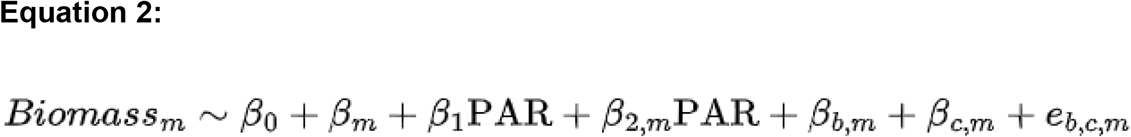

Then, we added light quality (µW/cm^2^/nm) to the baseline model, again assuming that each microgreen variety may have different sensitivity to light quality, where B, R, and FR represent the mean light intensity in the blue, red, and far-red light quality bands (µW/cm^2^/nm.), as specified in the previous section.

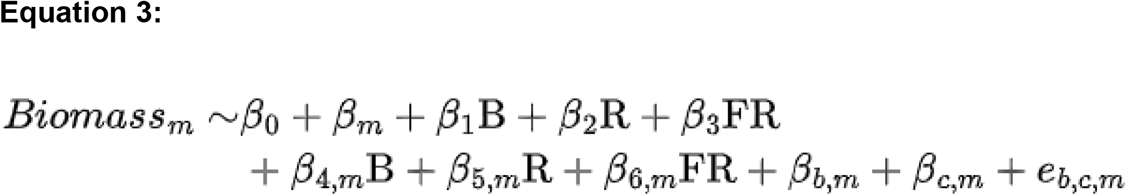

As many plant light sensors detect ratios of light quality, we also developed a model that incorporates light quality ratios, where R:B, R:FR, and FR:B represent the log10 ratios of red (R), blue (B), and far red (FR).

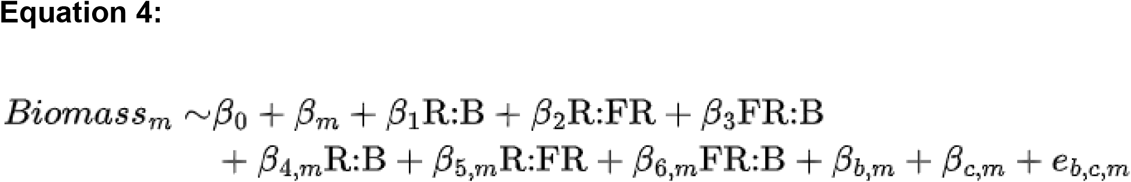

To compare the performance of these models we used an approach published by (22). The models with random effects were compared to the baseline model through separate ANOVAs, with all p-values compared to a 0.05 threshold after a Holm’s correction for multiple hypothesis testing (23). The Bayesian Information Criterion and Akaike Information Criterion were also assessed.

### Regularising general linear modelling

Equations 2-4 contain a large number of parameters compared to the number of observations, which introduces a risk of overfitting. We sought to use a well-established procedure to select a smaller set of parameters to include in our model. Specifically, a lasso regularisation procedure was employed to select a smaller subset of variables and prevent overfitting, using the glmnet package in R (24). Data was standardised prior to fitting, but the coefficients reported here are adjusted to be in the original scale. The initial model specification took the form:

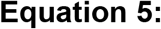

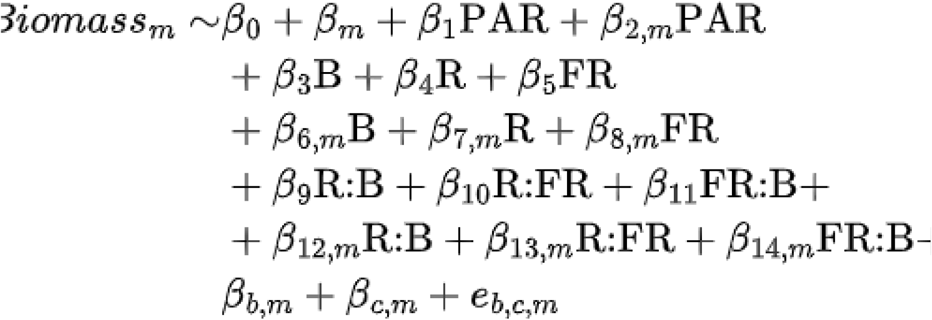

This model was evaluated using a two-level leave-one-out cross validation approach. For each of the 256 observations, a separate glmnet model was fit after leaving out one observation (i.e. 255 observations used to train the model). To select the lambda parameter of this model that would minimise the mean squared error, a further leave-one-out cross-validation strategy was deployed (using 254 observations for training the models for selecting the lambda parameter, by leaving out one of the 255 observations in the training set).

The output of this procedure was a set of 256 different models, each trained on 255 observations. For each model, we then predicted the biomass of the observation that was not used in training the model. This model fitting approach allows us to evaluate model performance in a way that is not affected by overfitting by testing our models on observations that were not used to train the model. Moreover, this approach provides us with a collection of 256 fitted values per coefficient, which enables us to evaluate how reliant our coefficient predictions are on the inclusion of any individual data point.

## Results

### Characterising the heterogeneity of light in a vertical farm

Our first aim was to quantify the level of light heterogeneity at plant level within the vertical farm. To do this we measured both PAR (**Figure 1A**) and spectral irradiance (**Figure 1B**) in 256 unique growing positions.

**Figure 1:**
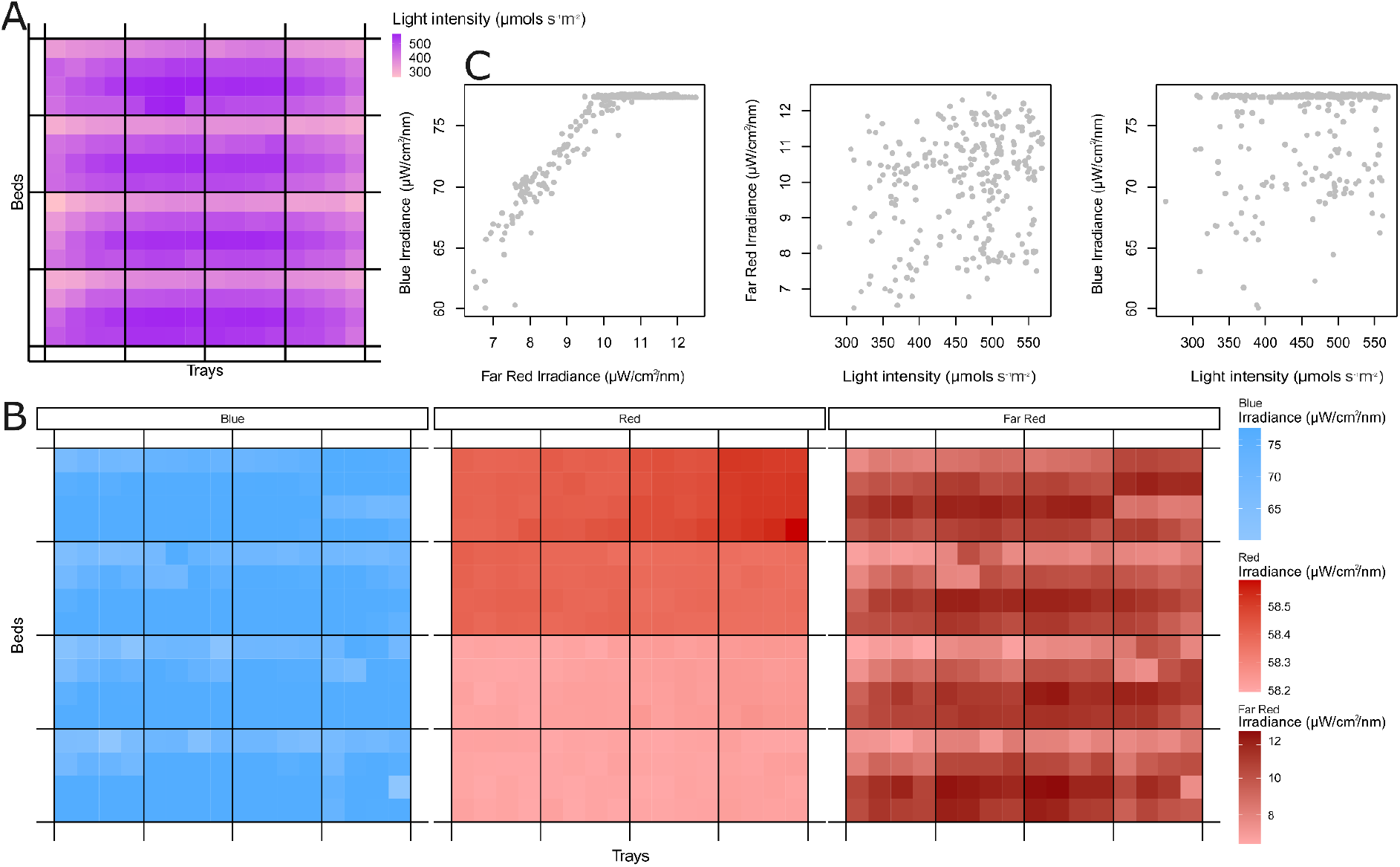
Heterogeneity of light quantity and quality within the vertical farm: (A) PAR readings and (B) average intensity within the following wavelength bands blue: 445-456nm, red: 653-668nm, far-red: 725-735nm. are shown across the farm. The beds are vertically stacked and each contains four trays with 16 positions per tray. Each square represents an area of size 83.7 cm^2^. Note that the plants grown along the edge of each tray are not included in this study and the light was not measured in these positions. (C) The relationship between blue and far-red irradiance and PAR readings, where each point represents a position/square in the images in (A) and (B).

The light intensity (PAR reading) was highest towards the front and centre of each bed (**Figure 1A**). A similar pattern of light intensity was observed for blue and far-red wavebands (**Figure 1B**), except for the rightmost trays in each bed. For red light, there was greater variation between beds than within beds, with the top two beds having higher levels than the bottom two (**Figure 1B**). Far-red and blue light intensities were highly correlated at low light settings, but blue light levels plateau at higher light intensities (**Figure 1C**). Although there was significant correlation between light intensity and blue and far-red light quality (P<0.001 for both), the Pearson correlation coefficient was low (R=0.22 and R=0.21, respectively). These results demonstrate that there is variation in both light quantity and light quality between and within growing beds in the vertical farm.

### Light quality is a predictor of biomass

In order to determine whether light quantity, light quality or ratios of light quality were predictors of biomass (the yield metric for microgreens), mixed effect models were constructed and compared to a baseline model that did not include any light-related parameters. Statistics for these comparisons are available in **Table 2**. The best-performing model was the light quality-based model (Equation 3), performing significantly better than the baseline model, even after Holm’s correction for multiple hypothesis testing (P<0.05). It also minimised the Akaike Information Criterion (AIC); however, this model performed poorly under the Bayesian Information Criterion (BIC) as BIC places a greater penalty on the number of parameters in the model. However, these results suggest that light quality can be used to predict biomass and that this is a better predictor of biomass than PAR readings alone.

**Table 2:** Summary of Mixed Effect Model Outcomes.

| Model | Equation | AIC | BIC | P | P adjust |
| --- | --- | --- | --- | --- | --- |
| baseline | Equation 1 | 1531.2 | 1556.0 | NA | NA |
| intensity | Equation 2 | 1529.7 | 1568.7 | 0.04987 | 0.09974 |
| quality | Equation 3 | 1526.0 | 1593.4 | 0.003724 | 0.01117 |
| ratios | Equation 4 | 1535.1 | 1588.2 | 0.1447 | 0.1447 |

This analysis was unable to determine whether combining information about light quantity and light quality ratios could further improve the model. In the next section, we will try to do so as well as attempt to minimise the complexity of the model.

### Variable selection highlights which aspects of light intensity and quality are most predictive of biomass

In order to determine which combination of explanatory variables was most predictive of biomass, we deployed a variable selection procedure via lasso regularisation (see Methods). Our simplified models were able to accurately predict the biomass of samples that were not used in training the dataset (see Figure 2A), with Pearson’s R=0.864 and P<0.05 (P=7.6e-78). Our predictions were also significantly correlated with the true biomass (P<0.05) for each of the individual varieties after Holm’s correction. Pearson’s R for kale 1 was 0.247, kale 2 was 0.487, radish was 0.597 and sunflower was 0.595. Notably, the variety for which our model performed the worst was kale 1, which was the lowest biomass variety and also the one with the least variance in biomass.

**Figure 2:**
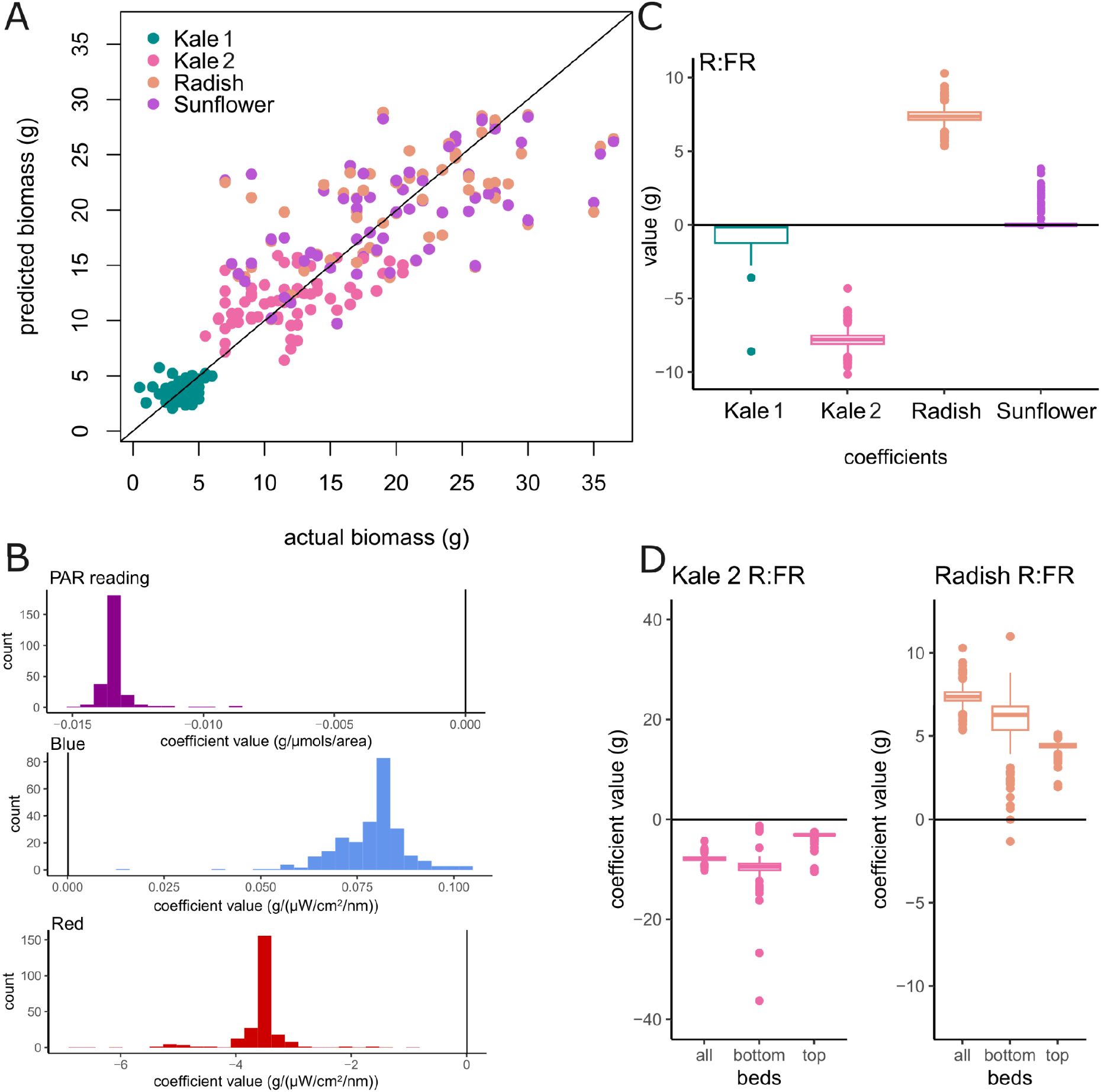
Analysis of lasso model coefficients. (A) Actual biomass at each position compared with predicted biomass when the lasso model was trained using all the positions except that one (leave-one-out cross validation, LOOCV). Biomass always refers to the total biomass (g) in each 83.7 cm^2^ sampling site. (B) Histogram of coefficient values for the LOOCV lasso models, for three different parameters (PAR reading, blue and red). (C) The variety-specific coefficients for the R:FR ratio across the LOOCV lasso models. (D) The variety-specific coefficients for kale (2) and radish, when different combinations of beds are used to train the model. Bottom refers to the bottom two beds, while top refers to the top two beds.

In further analysis of model coefficients, we decided to include all explanatory variables that were selected in at least 80% of the models (**Figure S4, Table S3**). Some light-related variables were associated with biomass in all microgreen varieties (coefficients from Equation 5): light intensity via PAR reading (β_1_) and blue (β_3_) and red (β_4_) light quality. Of these, red light was strongly negatively associated with biomass in all models, while blue light was positively associated (**Figure 2B**). Overall light intensity was slightly negatively correlated with biomass in all models (**Figure 2B**).

Although on their own, light quality ratios did not improve our ability to predict biomass compared to the baseline model, we found several light quality ratio variables that were selected in the combined model. However, these were all specific to individual varieties of microgreen. For instance, separate red:far-red ratio (referred to as R:FR in Figure 2 and below) coefficients were selected for each microgreen, with negative associations for both kale varieties and positive associations for radish and sunflower (**Figure 2C**). A full table of predicted coefficients for all models is available in **Table S2**.

### Variable prioritisation suggests R:FR ratio is a target area in kale and rocket production

To further validate the predicted associations between R:FR and yield in kale 2 and radish, we decided to re-fit the models independently using data from the top two beds and the bottom two beds. The top two beds and bottom two beds were found to contain relative differences in red light quality (**Figure 1B**). Additionally, they are\ likely to have unmeasured environmental differences, such as temperature or humidity gradients. Therefore, we wanted to confirm that R:FR was chosen by lasso regularisation in both cases and the effect of R:FR was consistent regardless of vertical positioning. The total number of variables selected was reduced, likely because we halved the number of observations used to train the model because only two of the four bays were used per analysis. As expected, the R:FR was selected as a key variable for radish and kale 2 in the top two beds, the bottom two beds, and in the combined model (**Figure 2D)**. Encouragingly, all three models predicted a negative coefficient of R:FR for Kale 2 and a positive coefficient of R:FR for Radish (**Figure 2D**). This suggests that the R:FR ratio is an appealing target for further spectral refinement, but that this needs to be performed in a variety-by-variety manner.

## Discussion

While there are some large vertical farming operations (25), many vertical farms (especially R&D vertical farms) are relatively compact, being situated in shipping containers (26), in distribution centres (27) or even in retail centres (28). For this reason, it may be impractical for these farms to iteratively perform light optimisation experiments, especially factorial designs that investigate the interactions between variables. Moreover, the knowledge obtained from optimising one vertical farm may not translate into optimal conditions in a different setting, as the light-dependency on yield may be dependent on the specific vegetable variety being grown (29) and other extrinsic conditions, like the temperature, humidity and growth medium (8). For instance, it has been found that red light can increase or decrease yield, depending on the wider growth context (30). It has been estimated that up to 85% of the carbon footprint of vertical farms comes from their high electricity demands (31), so it is paramount that vertical farms find the correct light recipes that effectively balance yield requirements and their carbon footprints.

For this reason, there is an immediate need to develop experimental designs that would prioritise the light qualities that are worth further investigation. We propose a three-phase workflow: (i) quantification of light heterogeneity within the facility, (ii) an observational experimental design and (iii) variable selection using lasso regularisation. The variables that are consistently selected and have high coefficients could be targeted for further investigation. Moreover, this workflow can suggest areas where heterogeneity in microenvironments could be most impacting the yield of the crops, areas warranting technological improvements to ensure greater environmental refinement. These could potentially be applied to other variables. Though we did not quantify temperature and humidity heterogeneity in this study, future experiments could include these variables.

As an initial demonstration of this workflow, our work highlights several key associations between light intensity, light quality and biomass. Firstly, heterogeneity in light quality within a vertical farm may have a greater impact on uniform crop yields than heterogeneity in light intensity. This is consistent with (32), which finds that light quality influences leafy green properties, even under uniform light intensity conditions. This suggests that it is important for small-scale vertical farms to track the heterogeneity of light quality in their facilities, instead of solely relying on PAR readings. Additionally, sensitivity to light ratios is likely to be variety-specific, which is consistent with several studies that have found different optimal light quality ratios were optimal for different varieties (33,34). This highlights how important it is to have ways of quickly screening light sensitivities in small-scale vertical farms, as a light recipe that works well for one variety is not guaranteed to produce optimal results for another.

The specific light sensitivities of the microgreens that we highlight are likely to be specific to the vertical farm facility and microgreen varieties we’ ve grown. Nevertheless, they indicate the kind of lessons that could be learnt from performing an observational study within a small-scale vertical farm. Our results have helped suggest light treatments that we can now test on our vertical farm to improve yield and reduce energy consumption.

## Supplementary Table Legends

Table S1: Spectral light readings

Table S2: Summary of PAR, blue, red, far-red light intensities, variety, and biomass, per position.

Table S3: Coefficients for Lasso

## Acknowledgements

We would like to acknowledge A. L. Tozer Limited for providing us with the kale seed for variety 2. We would also like to acknowledge the core R team that developed the version of R studio used in this study. Additionally we would like to thank Paul Scott, Jason Daff, Harry Stevens, Alison Fenwick, Jacob Woodward, Dave Grimshaw and Peter Smithson of the University of York Department of Biology horticulture department for their assistance and support with this work. The research presented in this paper was conducted while PS was with Vertically Urban; PS is currently affiliated with The UK Agri-Tech Centre. The authors would like to thank Vertically Urban for their support during the research period.

## Author contributions

WC and DE conceived the study. WC, DE, PS and KD designed the study. WC, AK and AT performed the experiments. WC, ER, GV and DE performed data analysis. WC and DE wrote and compiled the manuscript with contributions to the text from PS, KD, AK, AT, GV and ER.

