## Supplementary material for "Harnessing light heterogeneity to optimise controlled environment agriculture": Figures S1-S4

### Supplemental materials for “Harnessing light heterogeneity to optimise controlled environment agriculture”

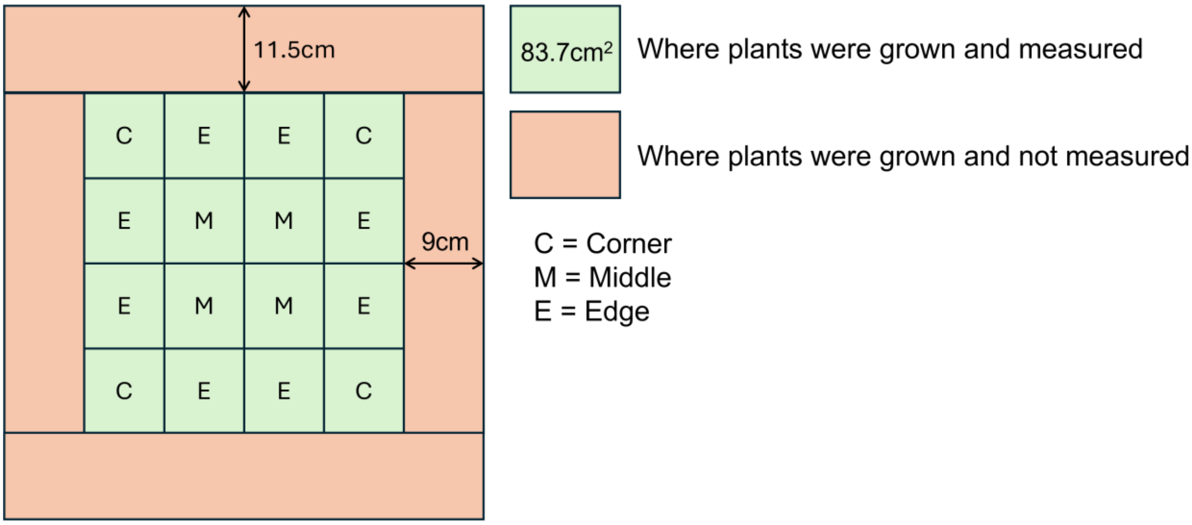


**Figure S1:** The layout of each tray.

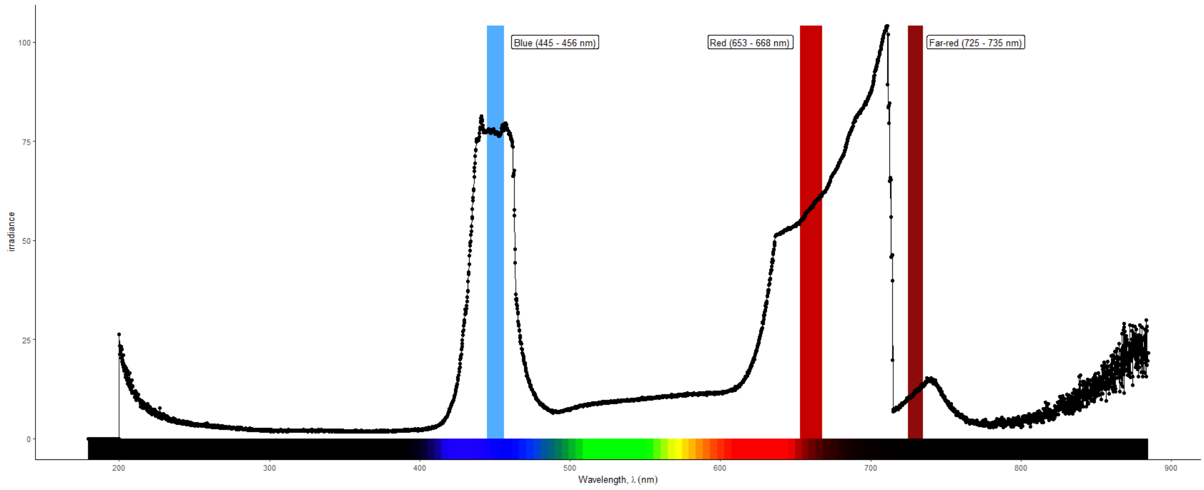


**Figure S2**: Example full light spectrum at plant level, with regions that we use to indicate blue, red, and far-red highlighted in coloured bars.

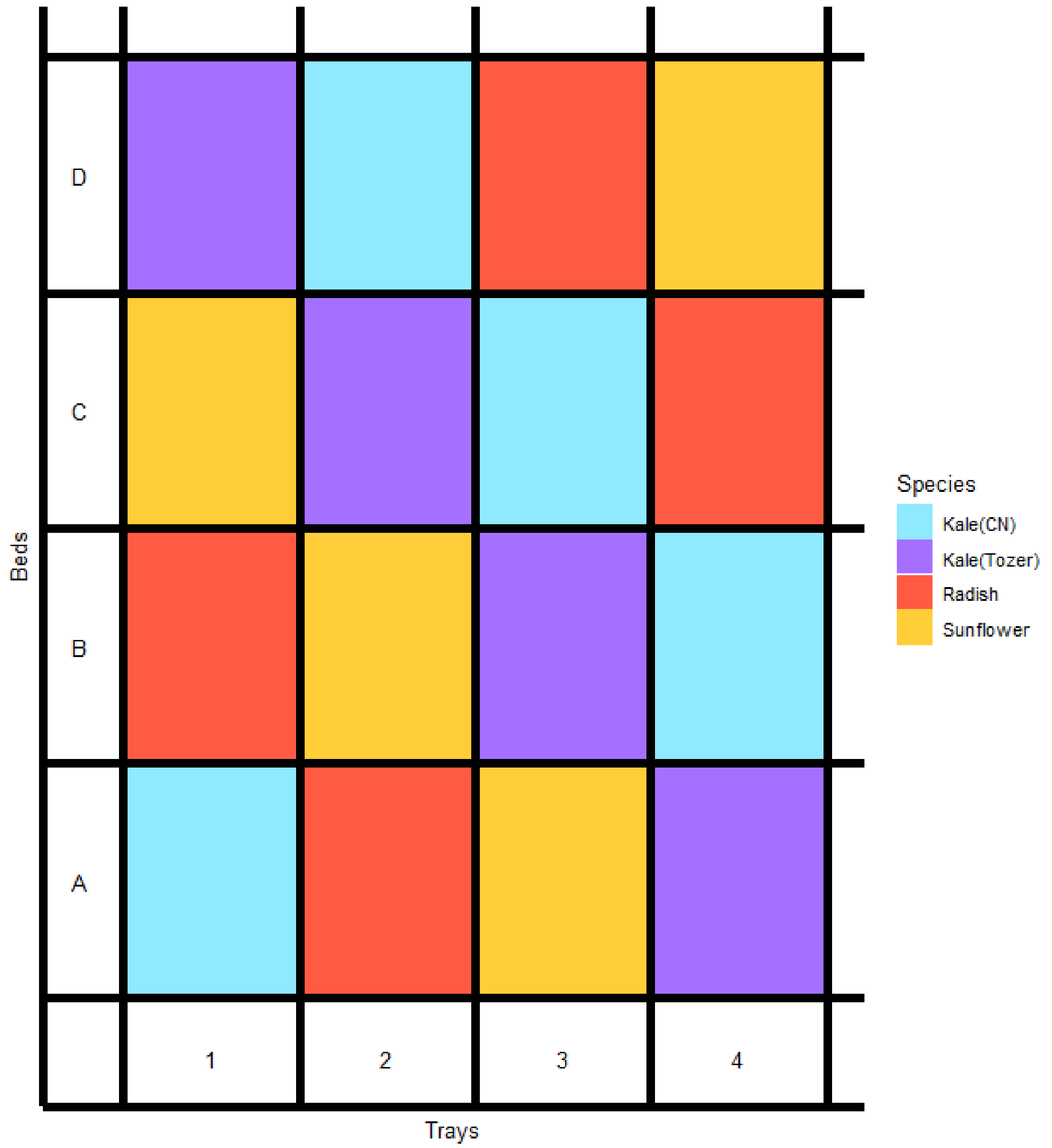


**Figure S3**: Arrangement of varieties in beds and trays.

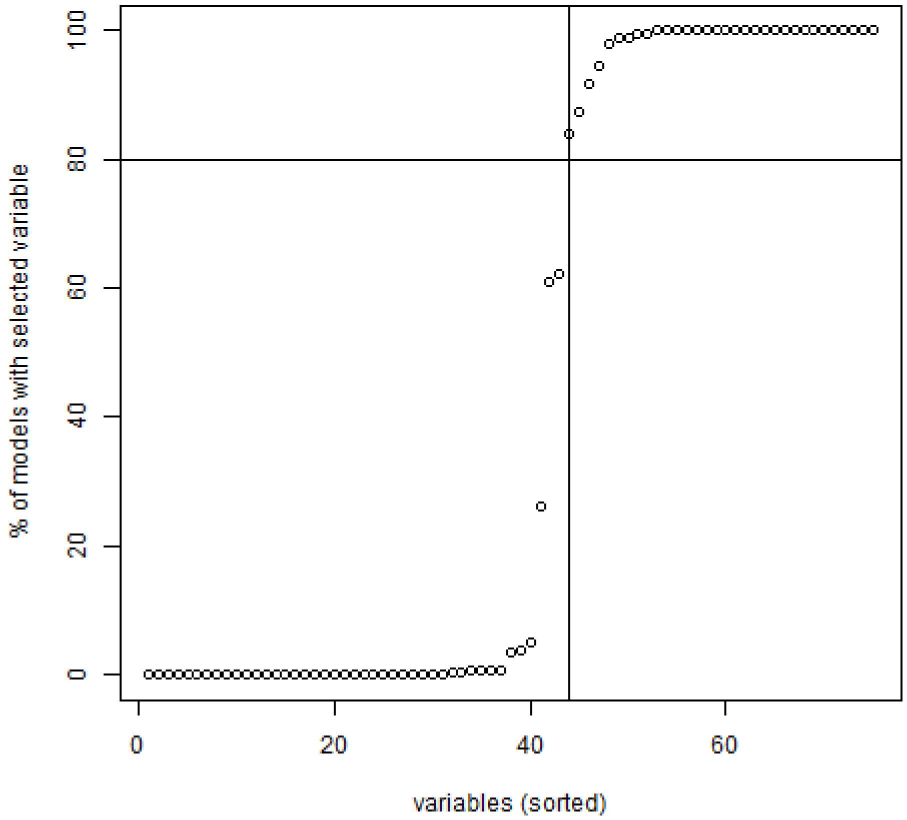


**Figure S4**: Proportion of models in which each variable was selected with non-zero coefficients.
